# The heritability of BMI varies across the range of BMI – a heritability curve analysis in a twin cohort

**DOI:** 10.1101/2022.01.06.475210

**Authors:** Francesca Azzolini, Geir D. Berentsen, Hans J. Skaug, Jacob v.B. Hjelmborg, Jaakko A. Kaprio

## Abstract

The heritability of traits such as body mass index (BMI), a measure of obesity, is generally estimated using family, twin, and increasingly by molecular genetic approaches. These studies generally assume that genetic effects are uniform across all trait values, yet there is emerging evidence that this may not always be the case. This paper analyzes twin data using a recently developed measure of heritability called the heritability curve. Under the assumption that trait values in twin pairs are governed by a flexible Gaussian mixture distribution, heritability curves may vary across trait values. The data consist of repeated measures of BMI on 1506 monozygotic (MZ) and 2843 like-sexed dizygotic (DZ) adult twin pairs, gathered from multiple surveys in older Finnish Twin Cohorts. The heritability curve and BMI value-specific MZ and DZ pairwise correlations were estimated, and these varied across the range of BMI. MZ correlations were highest at BMI values from 21 to 24, with a stronger decrease for women than for men at higher values. Models with additive and dominance effects fit best at low and high BMI values, while models with additive genetic and common environmental effects fit best in the normal range of BMI. Thus, we demonstrate that twin and molecular genetic studies need to consider how genetic effects vary across trait values. Such variation may reconcile findings of traits with high heritabilities and major differences in mean values between countries or over time.

## 1 Introduction

Twin and family studies of humans have provided evidence for genetic influences on anthropometric measures. One of the most studied phenotypes has been relative weight, the degree to which an individual is lean, of normal weight or has excess weight relative to their height. Body mass index (BMI), weight divided by height squared, is the most used measure in research due to the ease of its assessment and because among adults it is at most weakly correlated with height [1]. As excess weight is also associated with risk of cardiovascular and metabolic diseases [2], BMI is also in widespread clinical use.

Early meta-analyses on twins based on published summary data have shown that the heritability of BMI is generally high. The estimates based on twins are consistent with the patterns of resemblance of other first-degree relationships [3, 4]. These studies indicated that there is relatively little variation over age, but non-genetic familial influences seen in childhood and adolescence are largely absent in adults [5, 6]. By pooling individual data on height and weight from twin studies across the globe on over 140,000 twin pairs, [7] show that heritability of BMI decreases from young adulthood to old age, with relatively little differences by region or calendar time. Using the same resource with data from 87,782 twin pairs under the age of 20, [8] show that heritability of BMI was lowest in early childhood. Cross-sectional data from these twin and family studies, as well as from large molecular genetic studies [9] imply that genetic influences are fairly stable over the lifespan from early childhood onwards.

In contrast, longitudinal twin studies indicate that genetic influences do vary with age. Molecular genetic analyses suggest that different sets of genes act at different ages, both in childhood and among adults [10]. Twin analyses of children and adolescents show that as the individual develops and grows, there are novel genetic influences coming into play at different ages [11, 12]; these may reflect both changes in lean mass, such as muscle and bone growth, and in fat mass. Among adults, whose growth has ended, changes in weight result mainly from changes in body fat. Twin models indicate that genetic effects on weight change are poorly correlated with the stable component of BMI [13, 14]. Analyses of genetic risk scores at different ages support these results [10].

At different levels of BMI, the proportions of lean and fat mass differ on average, and hence it could be expected that genetic effects are not uniform across all BMI values. In an analysis of extreme leanness vs obesity, [15] found that the two traits were only partially correlated genetically (*rG* = 0.49). Using commingling analyses of BMI in MZ twin pairs, [16] found twin correlations to be lower in overweight and obese twin pairs. This restricts to a truncated upper-tail of the BMI distribution. The authors did not have DZ pairs to derive heritability estimates at different levels of obesity. Studies of the genetics of BMI across the whole spectrum of BMI are rare; a recent study uses parent-offspring and sibpair relationship and quantile regression to estimate heritability of BMI at various BMI values [17]. The study finds increasing heritability with increasing BMI values, a result seen with other measures of fatness but not height. However, using family relationships or only MZ pairs can be challenging to distinguish between genetic and non-genetic familial effects contributing to the estimated heritability.

Recently [18] extended the classical notion of heritability to that of a heritability curve, which allows the heritability to vary with the trait value. Using empirical data from the Finnish Twin Cohort, we demonstrate that there is variation in the contribution of genetic factors over the range of BMI values seen in a population sample.

## 2 Material and Methods

### 2.1 The dataset

The dataset we use in our analysis contains repeated BMI measurements on 4349 same sex twin pairs (1506 monozygotic and 2843 dizygotic) from the Finnish Twin Cohort [14], [19]. Each twin pair was asked to provide BMI measurements at different stages in life; for each pair we have up to 7 different values, at different ages and waves of data collections. Each such measurement is accompanied by the following information: the twin pair it belongs to, the wave number, the age at which the measurement is taken, the sex of the twins, and their zygosity. The data include information not only on weight at the current wave, but also recall of weight earlier in life.

Since the measurements were taken at different ages for different twin pairs, we use linear regression, separately on each individual, to obtain estimates of BMI at the reference age 35, which we denote with BMI_35_ (Figure 1). To reduce estimation uncertainty, we consider only twin pairs that have been measured three or more times. The resulting dataset contains 1493 monozygotic twin pairs (606 males and 887 females) and 2806 dizygotic twin pairs (1218 males and 1588 females). More details on the preprocess of the data can be found in the supplementary material.

**Figure 1:**
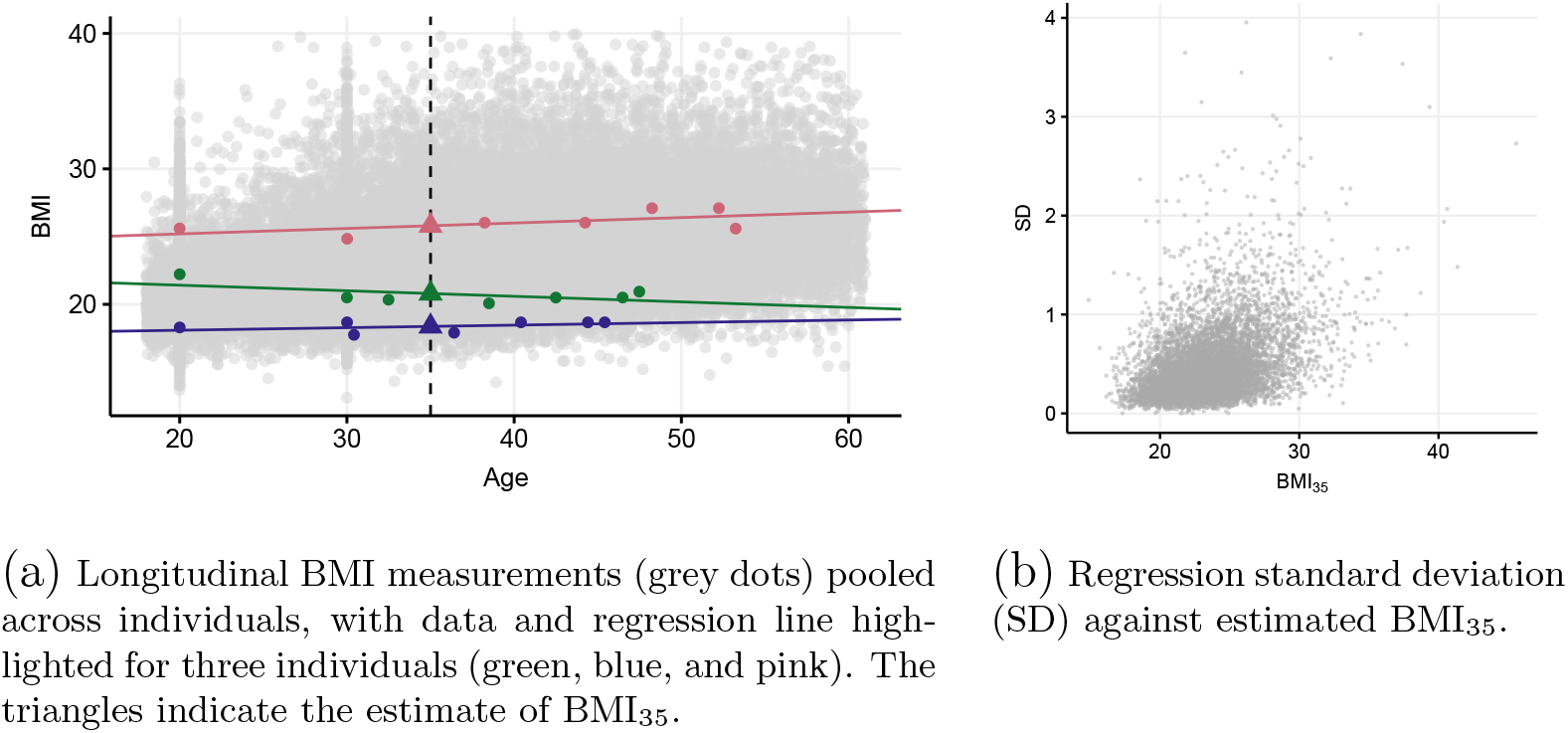
Regression analysis used to estimate BMI_35_ (BMI at age 35) for each of 8598 individuals in the study.

### 2.2 Statistical methods

#### 2.2.1 Heritability curve

In biometrical models, heritability is typically defined as the proportion of a trait variance attributed to genetic effects. Depending on the family structure of the data, the trait variance can be decomposed in several ways. The most commonly used biometrical model for twins is the ACE model, where it is assumed that the trait value can be decomposed into additive genetic effects (A), common (shared) environment (C), and residual (random) environment (E). The proportion of trait variance explained by component A is often referred to as narrow-sense heritability. Another frequently used model for twins is the ADE model, where the C component in the ACE model is replaced by dominant genetic effects (D). The proportion of trait variance explained by the components A and D combined is then referred to as the broad sense heritability [20].

Data on monozygotic and dizygotic twins provide contrasts from which the genetic variance can be separated from the environmental variance. For the ACE model, it is assumed that the amount of shared environment is the same for the two types of twins and that the amount of shared additive genetic effects is 100% and 50 % for mono- and dizygotic twins, respectively. If the correlation between the trait values of monozygotic twins is larger than the correlation between dizygotic twins, the difference is ascribed to the additive genetic effects alone, and the trait is heritable. The heritability can be quantified by comparing empirical correlations of monozygotic and dizygotic twins with the “expected” correlations implied by the model, and for the ACE model, this results in the well-known Falconer’s formula [21] for heritability:

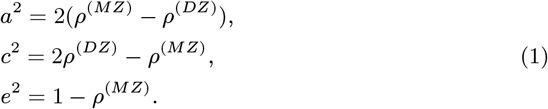

Here *a*^2^, *c*^2^ and *e*^2^ denote the proportions of the total variance explained by the components A, C, and E, respectively, while *ρ*^(*MZ*)^ and *ρ*^(*DZ*)^ denote the Pearson intraclass correlation of monozygotic and dizygotic twins, respectively. For the ADE model, the corresponding equations are given by

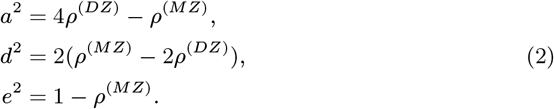

The derivation of equations 1 and 2 can be found in the supplementary material.

Recently, the classical notion of heritability has been extended to that of a heritability curve [18] assuming that trait values in pairs are governed by a Gaussian mixture distribution (see section 2.2.2). This allows the heritability to vary with the trait value, resulting in a curve *a*^2^ (*y*) that potentially varies for different trait values *y*. The heritability curves are derived based on the same type of variance decomposition as for ACE and ADE model, but conditionally on a given phenotypic value. In this way, the heritability curve measures the heritability as a function of the trait itself, and would not be expected to be constant over the whole phenotypic range. The conditioning on a phenotype value is done via local correlations curves [22]. Rather than comparing the ordinary Pearson correlation between phenotype values of monozygotic and dizygotic twins, we do the same type of comparison (e.g. using Falconer’s formula under the ACE model) using correlation curves *ρ_MZ_* (*y*) and *ρ_DZ_* (*y*), evaluated at different values of BMI. Note that this procedure also provides curves *c*^2^(*y*) (or *d*^2^(*y*) for the ADE model) and *e*^2^ (*y*) allowing the other components in the biometrical model to vary with the trait value as well. When there is no variation with trait value, the heritability curve reduces to the classical heritability coefficient.

#### 2.2.2 Gaussian mixtures

Classical heritability models assume a bivariate Gaussian distribution for pair of traits in twins, typically with a positive correlation. Under this assumption the heritability curve reduces to the classical heritability coefficient [18], and does not provide any additional insight. Bivariate Gaussian mixtures are a more flexible class of bivariate distributions and underlie the implementation of the heritability curve of [18]. A Gaussian mixture is a weighted sum of Gaussian kernels (Figure 2). The number *m* of Gaussian kernels is a data driven parameter, and for this purpose we use the BIC criterion [18]. Note that with *m* = 1 the mixture reduces to an ordinary bivariate Gaussian distribution. Each bivariate Gaussian kernel has three parameters: mean, variance and correlation, i.e. mean and variance are assumed identical across the twin individuals, yielding in total 3 times *m* unknown parameters. Monozygotic and dizygotic twins are allowed to have different correlations, which introduces *m* additional correlation parameters. Finally, there are *m* weight parameters, but due to a sum-to-one constraint, only *m* – 1 of these need to be estimated. The total of *Q* = 5*m* – 1 parameters are estimated by maximum likelihood [18]. Formulae exist for the marginal mean, variance and correlation, referred to as “global”, in terms of the kernel-specific parameters.

**Figure 2:**
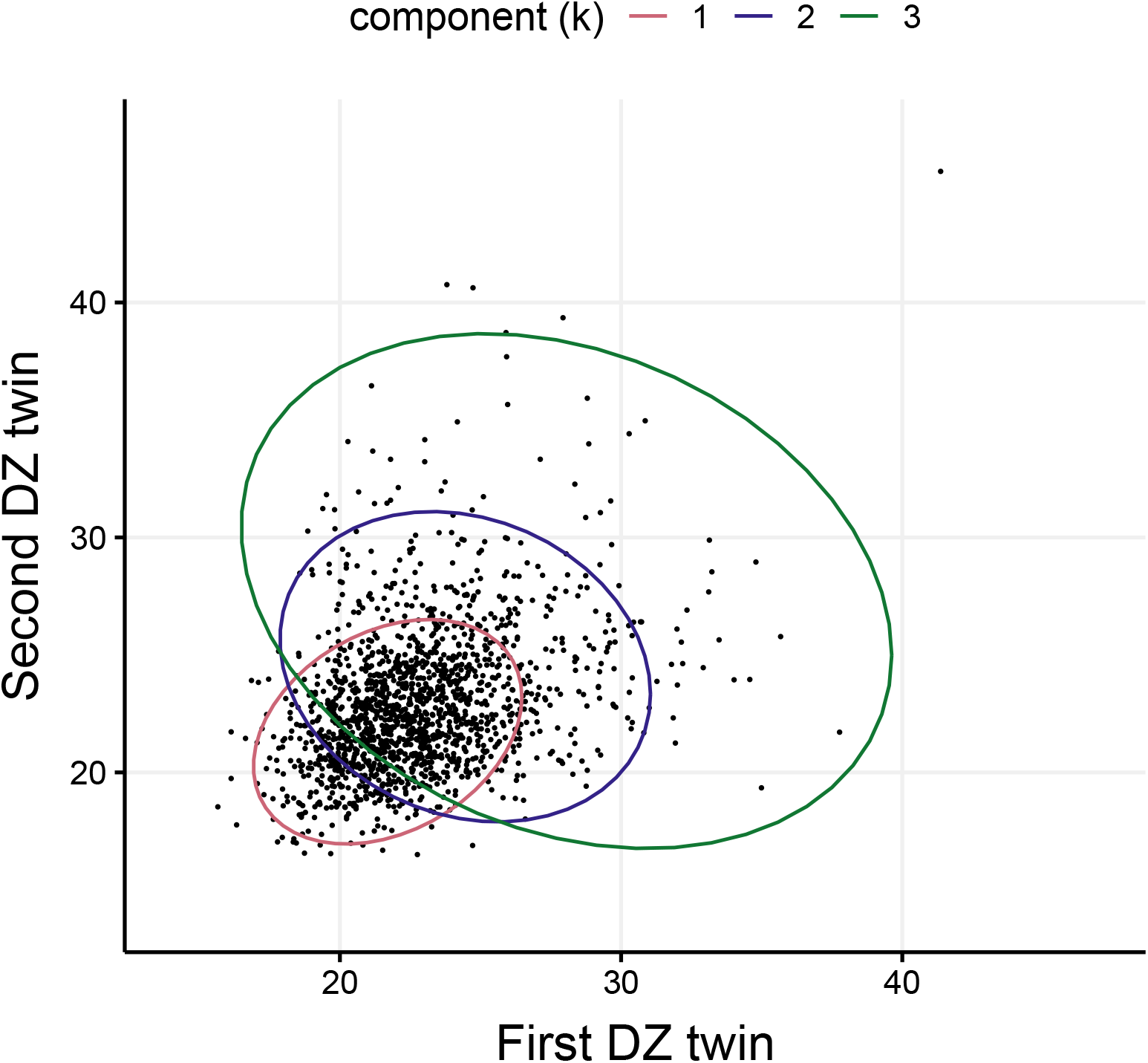
BMI_35_ for pairs of female dizygotic twins, obtained by regression analysis. The three ellipses represent 0.95 probability regions for the three Gaussian kernels of the fitted mixture distribution. The parameter values associated with each kernel can be found in Table 2.

Covariates can be introduced into both the mean, variance, or covariance part of the model. In our study sex is the only covariate, and we consider three different configurations of the model:

1. “Stratified” in which fully separate Gaussian mixtures are fitted for males and females, except that they are constrained to have the same value of *m*,
2. “Mean” in which only the mean is sex specific. The mean of each Gaussian kernel for males is right-shifted by the same amount from females.
3. “Mean+covariance” in which *m* mean and correlation parameters are sex specific, but the variance is assumed equal by sex.

A mathematical description of the model is provided in supplementary material. The supplement also contains details about how the parameters of the Gaussian mixture were estimated from data using the software package TMB [23].

#### 2.2.3 Biometrical model selection

The choice between the ACE and ADE model relies traditionally on the relationship between the (empirical) Pearson correlations *ρ*^(*MZ*)^ and *ρ*^(*DZ*)^. More specifically, the sign of the quantity 2*ρ*^(*DZ*)^ – *ρ*^(*MZ*)^ indicates which model is more appropriate. Under the ACE model, this quantity corresponds to the proportion of variance explained by the shared environment, *c*^2^. By contrasting equations (1) and (2) we obtain the relationship *d*^2^ = −2*c*^2^ with the proportion of variance explained by the dominant genetic effects, *d*^2^. Consequently, if 2*ρ*^(*DZ*)^ – *ρ*^(*MZ*)^ < 0, i.e. *c*^2^ is negative, the ACE model is (most likely) misspecified. Vice versa, if 2*ρ*^(*DZ*)^ – *ρ*^(*MZ*)^ > 0, *d*^2^ is negative and the ADE model is misspecified.

In the context of correlation curves, the ACE model may be suitable in some part of the phenotypic range, while the ADE model may be more appropriate in the remaining part. Adopting the above procedure, we can then switch between ACE and ADE model according to the sign of the quantity 2*ρ*^(*DZ*)^(*y*) – *ρ*^(*MZ*)^(*y*); the ACE model is preferred when 2*ρ*^(*DZ*)^(*y*) – *ρ*^(*MZ*)^(*y*) *>* 0, and the ADE model otherwise [18].

## 3 Results

The longitudinal BMI measurements used in the regression analysis are shown in Figure 1a. The distribution of uncertainties in fitted BMI_35_ is mostly confined to the interval from 0 to 2, but some twin pairs have higher uncertainty (Figure 1b). The bivariate distribution of BMI_35_ within twins deviates from normality for DZ females and is having a pear shape with less association for large BMI values (Figure 2). The same is true for males, and to a lesser extent for MZ twins (Supplementary material). The mixture model can accommodate this pear shape, by using *m* = 3 individual Gaussian components (Figure 2). The fact that the mixture distribution fits data much better than a bivariate Gaussian distribution (*m* = 1) is clear from a comparison of BIC values (Table 1).

**Table 1:**
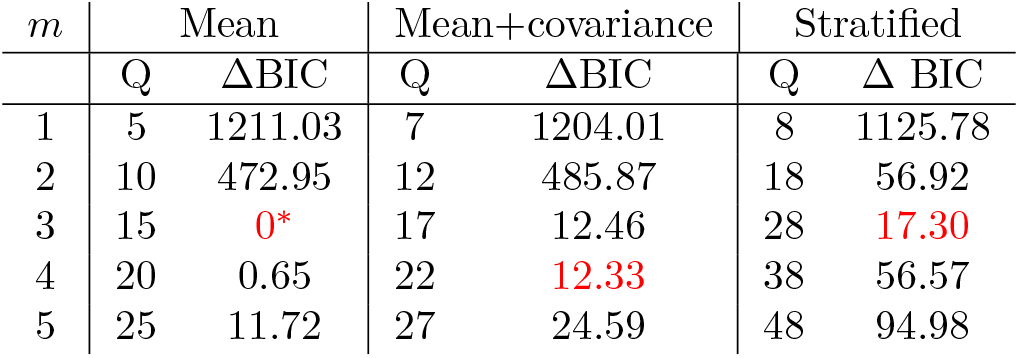
Model selection by the BIC criterion among three candidate models for sex effect (columns) and the number of mixture components (*m*). *Q* represents the number of parameter estimated, and ΔBIC shows BIC relative to the best fitting model (*) across the table. Color red highlights the model with lowest BIC within each column.

The best fitting covariate model is that in which sex affects only the mean of the response (Table 1). However, the difference in terms of BIC between this model and the stratified model or the model with a sex effect also in the correlation structure is not large. The latter two are both more flexible in their ability to fit the distributional shape of data but are being penalized by the BIC criterion for having more parameters than the selected model. The selected model has *m* = 3 mixture components. The BIC values in Table 1 are for the entire dataset (male/female and MZ/DZ). In the stratified model, this amounts to adding the BIC values computed separately for males and females. In an additional analysis, where males and females were allowed to have a different value of *m* it was found that the best fitting values were *m* = 2 for males and *m* = 3 for females, but the total BIC was not lower than the selected model in Table 1. A more in depth analysis of the different models can be found in the supplementary material.

The parameter estimates for the mixture model (Table 2; column “Global”) show that mean BMI is higher for males than for females by *μ*_male_ – *μ*_female_ = 1.86 units. The global correlation is expectedly stronger in MZ twins than in DZ twins (*ρ*_MZ_ = 0.70 versus *ρ*_DZ_ = 0.34). By constraint of the chosen model (Table 1), correlations are the same for males and females. For the three individual Gaussian components of the mixture, components *k* = 2, 3 have a negative correlation for DZ twins, which is contributing to the pear shape of the overall mixture (Figure 2). Component *k* = 3 has the largest standard deviation (*σ*) and represents 100 × *p*_3_ = 4% of the data (Table 2). This part of the data includes twin pairs which may be classified as outliers in the sense of having a strong negative association in BMI across the twins (Figure 2).

**Table 2:** Parameter estimates for the chosen *m* = 3 component Gaussian mixture assuming a sex effect in the mean (*μ*). Additional parameters are standard deviation (*σ*), monozygotic and dizygotic correlation (*ρ*), and mixture weights *p*. Each column (*k* = 1, 2, 3) corresponds to different mixture components. The final column refers the parameter value for the mixture as a whole.

| Parameters | $k = 1$ | $k = 2$ | $k = 3$ | Global |
| --- | --- | --- | --- | --- |
| $\mu_{\text{male}}$ | 23.57 | 26.36 | 29.82 | 24.55 |
| $\mu_{\text{female}}$ | 21.70 | 24.49 | 27.96 | 22.68 |
| $\sigma$ | 1.94 | 2.72 | 4.67 | 2.84 |
| $\rho_{\text{MZ}}$ | 0.74 | 0.34 | 0.38 | 0.69 |
| $\rho_{\text{DZ}}$ | 0.31 | -0.19 | -0.22 | 0.36 |
| $p$ | 0.70 | 0.26 | 0.04 | |

By construction of the selected model, the shape of the correlation curve for males is identical to that for females, but is right-shifted by an amount *μ_male_* – *μ_female_* = 1.87 (Figure 3). While this might seem like a strong restriction, it is worth noting that our model is preferred by the BIC criterion over the two other models in Table 1, which both allow for more flexibility in the correlation curves. Twin [7] and molecular genetic studies of BMI [9] have shown very little evidence of sex-specific genetic variance or genes expressed only in one sex. In contrast, the distribution of fat differs between men and women, and is affected by genetic factors.

**Figure 3:**
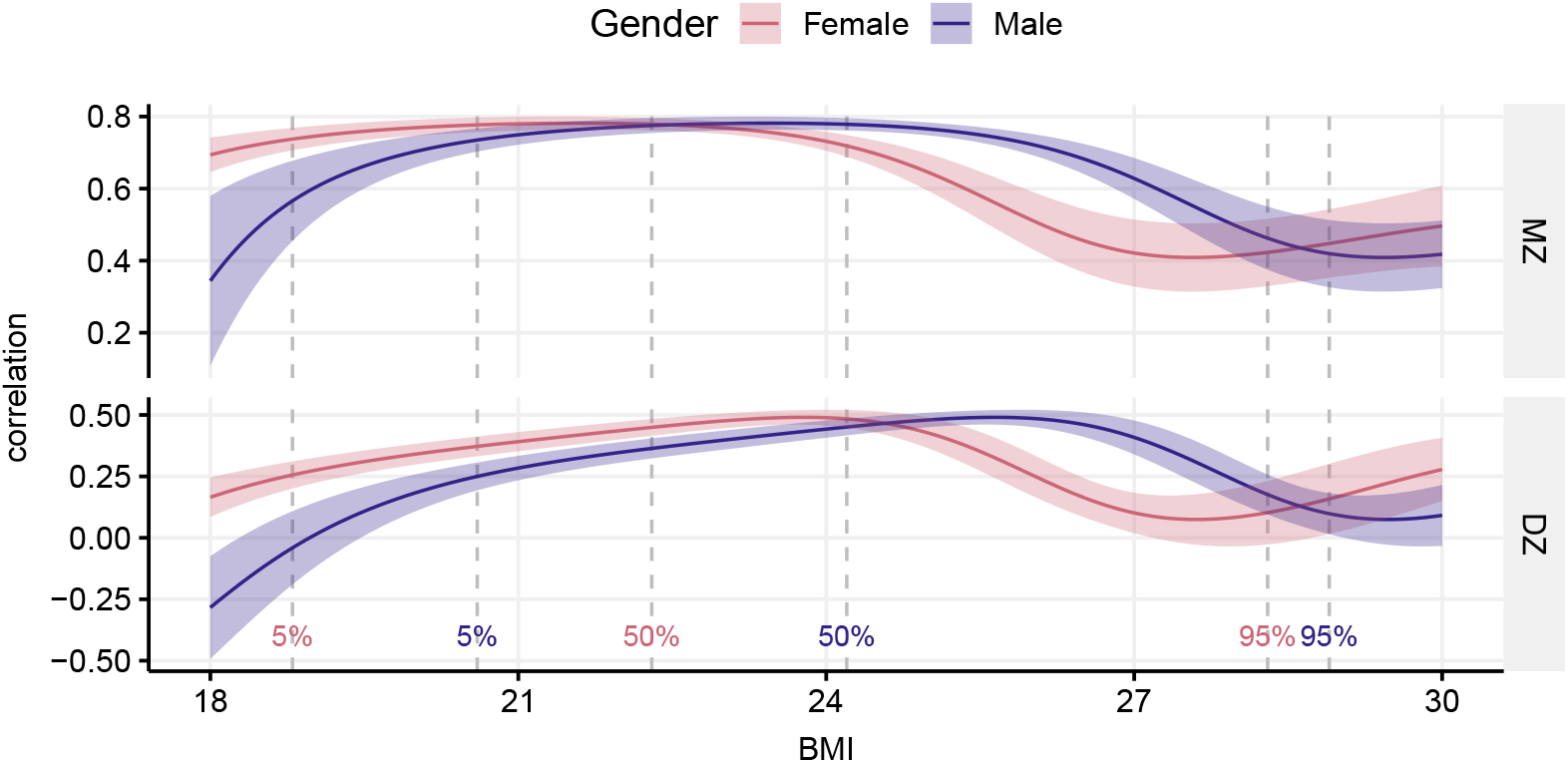
Correlation curves for male and female data for monozygotic (top) and dizygotic (bottom) twins, with pointwise 95% confidence bands (shaded regions). Vertical dashed lines show empirical quantiles (not model dependent) separately by sex, but pooled over MZ and DZ twins.

We observe a drop in the correlation curve for high BMI in both monozygotic and dizygotic twins (Figure 3). The correlations increase before dropping, but this pattern is more noticeable in dizygotic twins (it increases up to a BMI value of about 24 for female data and 26 for male data).

The property that the male correlation curve is, by construction, just a right-shifted version of the female one carries over to the heritability curves. Hence, we only discuss female heritability in the following. Figure 4 displays curves obtained using both ACE and ADE genetic models. The panel labeled “common” contains the dominant genetic component (which appears in the ADE model) and the shared environment (which appears in the ACE model). The curve for the residual environment is the same in both models.

**Figure 4:**
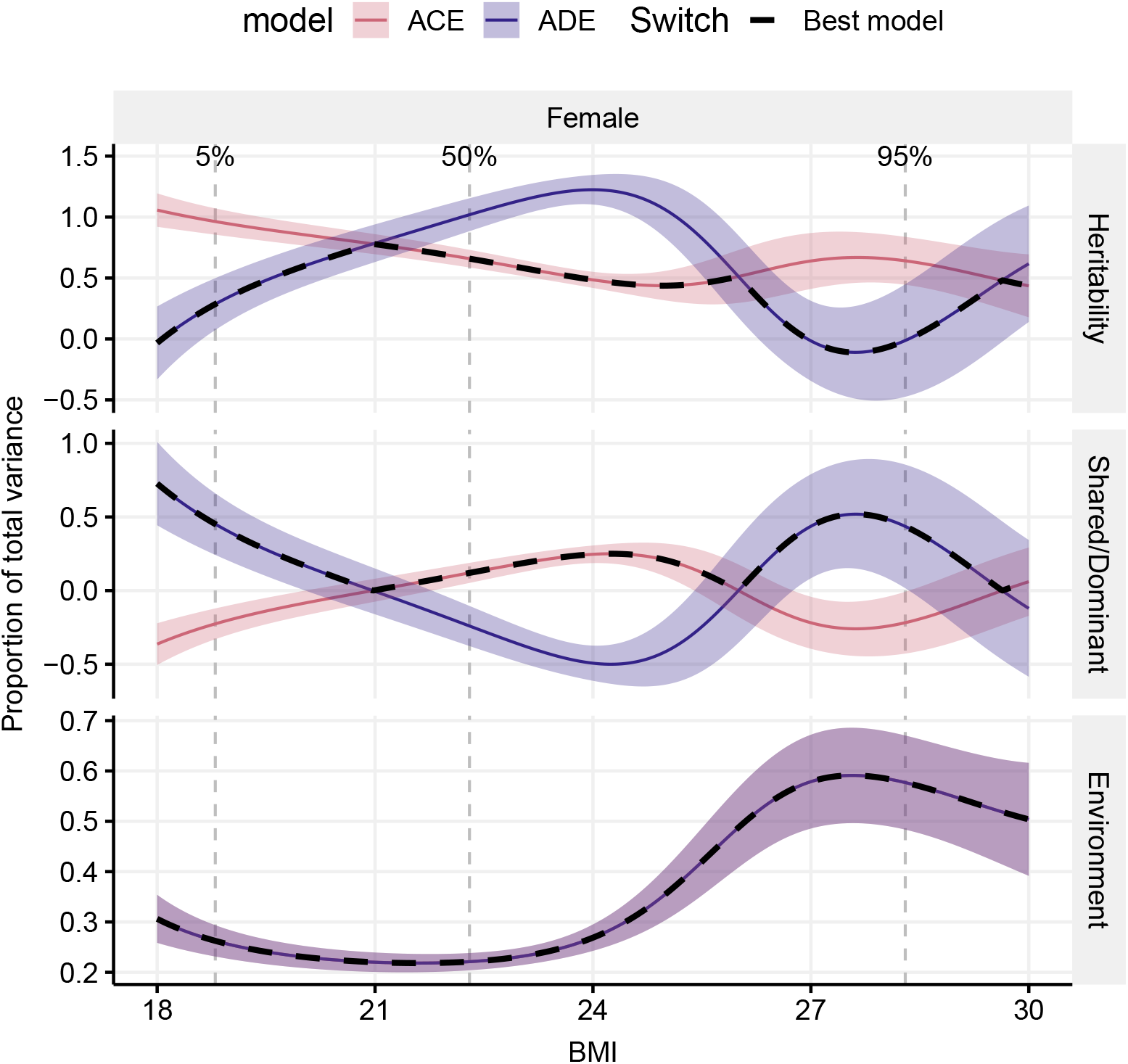
Decomposition of total variance into genetic and envirolmental effects by BMI, sex, and genetic model (ACE/ADE). Shaded areas indicate 95% pointwise confidence bands. The environmental part is identical for the ACE and ADE models, indicated by the pink color in the lower part. The dashed black line represents the combination of the ACE and ADE models.

The Pearson correlations are *ρ*^(*MZ*)^ = 0.74 for monozygotic twins and *ρ*^(*DZ*)^ = 0.42 for dizygotic twins. The quantity 2*ρ*^(*DZ*)^ – *ρ*^(*MZ*)^ is positive; hence, if we opt to use one single model for the whole dataset range, the ACE model is most appropriate. However, as discussed in Section 2.2.3 we can instead switch between ACE and ADE model depending on the sign of the quantity 2*ρ*^(*DZ*)^(*y*) – *ρ*^(*MZ*)^ (*y*). The dashed line in Figure 4 indicates the preferred (combined) model. We define the “mid range” of BMI values (21–27) as the region where the ACE model is preferred and “low/high range” BMI values as the region where the ADE model is preferred.

The mid range BMI values are governed by additive genetic effects (A) and to some extent the shared environment (C). The residual environment (E) plays a larger role for the upper mid-range BMI values. The heritability curve *α*^2^ steadily decreases while the BMI values increase, starting from its highest value of 0.78 for the BMI value 21. The shared environment curve *c*^2^, instead, displays a convex shape, increasing together with the BMI up until it reaches its maximum value of 0.25 around a BMI value of 24 and decreasing after. By construction, *c*^2^ is zero at both extremes.

Low BMI values are overall governed by the genetic effects A and D. Dominant genetic effects (D) play a more pronounced role as the BMI values decrease, with its maximum value (0.73) reached at BMI value of 18, while the opposite trend can be seen with the additive heritability curve *a*^2^. We also see a slight increase in environmental effects (E) as the BMI values decrease.

High BMI values are increasingly governed by environmental effects (E) (for a maximum value of 0.60) while broad sense heritability is decreasing (Figure 4). Interestingly, a change in type of genetic action is suggested by the curves; while additive genetic effect, the A component, is the suggested action for BMI values in the normal range, dominant genetic and weak epistatic effects (the D component) tend to govern the upper part of the BMI scale effects (D) ((for a maximum value of 0.52 around a BMI value of 28). The basis for this suggestion follows from decreasing within-pair correlation in BMI among MZ pairs with increasing BMI and similar decreasing within-pair correlation among DZ pairs, however with larger (or faster) decrease for the DZ pairs with increasing BMI. Implications of this suggested change in mode of genetic influence is discussed below.

## 4 Discussion

In the present analysis, we demonstrate that multiple mixture models account for the pairwise relationship of BMI rather than a single bivariate distribution assumed in prior analyses of twin data of BMI. Further, the 95% probability regions of the kernels of the distributions are shaped differently as seen in Figure 2. The majority of pairs are in a symmetric circular distribution, while the remainder are in distributions indicating greater within pair differences, possibly due to greater than average genetic differences and/or specific environmental triggers affecting body weight development more in one twin than in the other. Such pairs discordant for BMI have proved very informative for the study of causes and consequences of obesity [24], [25]. The existence of several types of bivariate distributions suggests that a single bivariate normal model of multifactorial inheritance with a polygenic component is not sufficient to account for the complexity of interplay of genetic and environmental factors in BMI, even though GWA studies have been highly successful in identifying hundreds of BMI-associated genes and accounting for about a fifth of the variance in BMI [9]. On the other hand, rare Mendelian variants and various obesity-related syndromes account for a relatively small proportion of variance in BMI [26].

When we consider the resulting heritability curves, and associated curves of MZ and DZ correlations by level of BMI, we observe very high estimates of the contribution of genetic factors in the region of what is generally termed normal BMI [27]. As BMI comprises both lean (muscle, organs and bone) as well as fat mass, our results are consistent with the notion that in the absence of excess fat, body build is highly genetic determined. As BMI increases, the proportion of weight accounted for by fat mass increases, and the contribution of genetic variation decreases. This is consistent also with the rapid increase in obesity in global populations [28] being due to environmental factors rather than changes in the gene pool over the past decades.

In the ADE model, the genetic effect is split into an additive genetic component (A) and a dominant genetic component (D). We can compare the curve *a*^2^(*y*)*_ACE_*, computed using the ACE model, with the sum *a*^2^(*y*)*_ADE_* + *d*^2^(*y*), as they both represent the total heritability of the model (the latter assuming independence of A and D). Hence, even though a first look at Figure 4 may suggest a contradiction between the heritability estimate in the ACE and ADE models, we should not compare *a*^2^ (*y*)*_ACE_* and *a*^2^(*y*)*_ADE_*. Instead, we can study the behavior of the separate components of the total heritability. In particular, the dominant genetic component plays a larger role on the tails, while the additive genetic component has a larger effect in the middle of the data range.

Noteworthy, as can be derived from the biometric model, the D component may reflect some evidence for epistatis besides the dominant effects of variants. Hence the heritability curves may shed light on values for which such action may take place.

For BMI, it is biologically plausible that the genetic and environmental components vary over the range of BMI. For example, very large or small values of BMI could be caused by “sporadic” environmental factors such as accidents or by rare genetic mutations whereas the medium phenotype variation may be dominated by multiple common genetic factors. The heritability curves then provide insights to an evolutionary normal spectrum of BMI of which magnitude of genetic variants is observed. Expectedly, genetic action on BMI for values outside the normal spectrum may stem from different localizations of variants governing different mechanisms. Hence the curves relate to some combination of genotypic, environmental and epigenetic interactions, the broadsense heritability and it becomes important to study how curves may change given observed genetic variants which will be a perspective for further studies of correlation curves.

In this paper, we use a Gaussian mixture distribution to fit the data. To test the soundness of this assumption, we fitted a non-parametric correlation curve and showed that it returns similar results to Figure 3 everywhere except for low BMI values for female dizygotic data, where it estimates a higher correlation. See supplementary material for more details.

## Data Availability

The FTC data is not publicly available due to the restrictions of informed consent. However, the FTC data is available through the Institute for Molecular Medicine Finland (FIMM) Data Access Committee (DAC) for authorized researchers who have IRB/ethics approval and an institutionally approved study plan. To ensure the protection of privacy and compliance with national data protection legislation, a data use/transfer agreement is needed, the content and specific clauses of which will depend on the nature of the requested data.

## Author Contributions

FA, GDB, HJS and JBH contributed to the conception and design of the work. JBH and JAK provided the data material. FA carried out all statistical analyses. FA drafted the manuscript, except for sections Introduction and Discussion which were drafted by JAK. All authors participated in finalizing the manuscript, and gave final approval and agreed to be accountable for all aspects of work ensuring integrity and accuracy.

