## Supplementary material for "The heritability of BMI varies across the range of BMI – a heritability curve analysis in a twin cohort"

#### 1 Data preprocessing by linear regression

The purpose of this supplementary is to provide additional information about the analysis in "The heritability of BMI varies across the range of BMI – a heritability curve analysis in a twin cohort" (Azzolini, Berentsen, Skaug, Hjelmborg, Kaprio), which is referred to as "main text".

The main text studies the BMI of twins using the Finnish Twin Cohort [1]. The relevant variables to the analysis are listed in Table 1.

Since the measurements are taken at different ages for different twins, we need to preprocess the data to obtain a set of comparable values. For each twin, we interpolate in up to seven BMI measurements from the different waves using simple linear regression to obtain BMI estimates at age 35 (approximately the average age in the dataset).

| Name | Description | Values |
|---|---|---|
| BMI | BMI measurements | $[13.11, 29.92]$ |
| Age | Age of the twin pair at measurement | $[18, 61]$ |
| Sex | Sex of the pair (only same-sex pairs are included) | 1 (Male), 2 (Female) |
| Zygoty | Zygosity of the twin pair | 1 (MZ), 2 (DZ) |
| Wave | Different measurements of the same twin pair | from 1 to 7 |
| Tvparnr | ID number of the twin pair | from 1 to 7639 |
| Twinnumber | Different twins in the same pair | 1,2 |
| Twinnid | summary of both Tvparnr and Twinnumber | from 11 to 76392 |

Table 1: List and description of the variables included in the dataset.

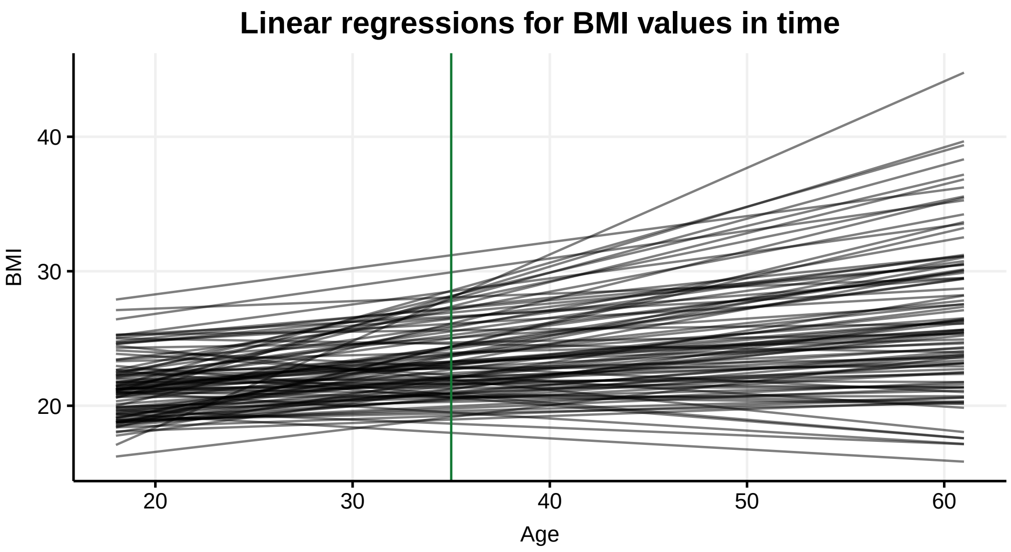


Figure 1: Regression lines fitted to BMI-at-age for 100 randomly selected individuals. The vertical green line is at age 35.

We use the program R [2] for the computations. We show a simplified version of the code in appendix A.

Figure 1 shows the regression lines of 100 randomly selected individuals from the real data, obtained through the process explained above.

#### 2 Derivation of classical heritability formulas

In this section we show how to derive equations (2.1) and (2.2) in the main text.

Let $Y_{ij}$ be the trait value of twin $j$ ($j = 1, 2$) in twin-pair $i$, and let $\rho^{(MZ)}$ and $\rho^{(DZ)}$ be the Pearson correlations $\mathrm{cor}(Y_{i1}, Y_{i2})$ for monozygotic and dizygotic twins, respectively. Consider the mixed-effect model [3]

$$Y_{ij} = \mu + \beta^t x_{ij} + A_{ij} + C_{ij} + D_{ij} + E_{ij}, \tag{1}$$

where $A_{ij}$, $C_{ij}$, $D_{ij}$ and $E_{ij}$ are mutually independent and follow a normal distribution with mean 0 and variances $\sigma_A^2$, $\sigma_C^2$, $\sigma_D^2$ and $\sigma_E^2$, respectively. The total variance of $Y_{ij}$ is $\sigma^2 = Var(Y_{ij}) = \sigma_A^2 + \sigma_C^2 + \sigma_D^2 + \sigma_E^2$. Note that this model assumes no gene-environment interaction. We define $a^2 = \sigma_A^2/\sigma^2$, $c^2 = \sigma_C^2/\sigma^2$, $d^2 = \sigma_D^2/\sigma^2$, and $e^2 = \sigma_E^2/\sigma^2$. By definition,

$$a^2 + c^2 + d^2 + e^2 = 1, \tag{2}$$

i.e. the contributions from all components sum to one.

The ACE and ADE models assume, respectively, that dominant genetic effects and shared environment do not affect the trait in study; in other words, $d^2$ and $c^2$ are assumed to be zero.

Mono- and dizygotic twins share 100% and 50% of the additive genetic effects, respectively. In formula, we write $\mathrm{cor}(A_{i1}^{MZ}, A_{i2}^{MZ}) = 1$ and $\mathrm{cor}(A_{i1}^{DZ}, A_{i2}^{DZ}) = 1/2$.

Monozygotic and dizygotic twins alike share the totality of the common environment, hence we make the common assumption $\mathrm{cor}(C_{i1}, C_{i2}) = 1$. The dominant genetic component, instead, affect monozygotic and dizygotic twins differently; in particular, $\mathrm{cor}(D_{i1}^{MZ}, D_{i2}^{MZ}) = 1$ and $\mathrm{cor}(D_{i1}^{DZ}, D_{i2}^{DZ}) = 1/4$.

Traditional twin models utilize only the $\rho^{(MZ)}$ and $\rho^{(DZ)}$ phenotype correlations, which for the ACE model are

$$\begin{aligned} \rho^{(MZ)} &= \quad a^2 + c^2, \\ \rho^{(DZ)} &= \frac{1}{2}a^2 + c^2, \end{aligned} \tag{3}$$

while for the ADE model are given as

$$\begin{aligned} \rho^{(MZ)} &= a^2 + d^2, \\ \rho^{(DZ)} &= \frac{1}{2}a^2 + \frac{1}{4}d^2. \end{aligned} \tag{4}$$

Equations (2.1) and (2.2) in the main text can be easily derived from 2, 3, and 4.

### 3 Bivariate Gaussian mixtures

[4] describe in depth the advantages of using Gaussian mixture models as underlying distributions when constructing correlation curves. We summarize below the model that we use in this analysis, but we refer the interested reader to [4] for more details.

The probability density of a $m$-component Gaussian mixture for a twin phenotype $\boldsymbol{y} = (y_1, y_2)$ is

$$\sum_{k=1}^{m} p_k \phi_2(\boldsymbol{y}; \boldsymbol{\mu}_k, \boldsymbol{\Sigma}_k)\,. \tag{5}$$

The parameters $p_1, \ldots, p_m$ are non-negative values satisfying $\sum_{k=1}^{m} p_k = 1$, and $\phi_2(\boldsymbol{y}; \boldsymbol{\mu}, \boldsymbol{\Sigma})$ denotes a bivariate normal density, with mean vector $\boldsymbol{\mu}$ and covariance matrix $\boldsymbol{\Sigma}$.

To guarantee symmetry between the two twins, we impose some conditions on the mean vector and the covariance matrix:

$$\boldsymbol{\mu}_k = (\mu_k, \mu_k), \quad \boldsymbol{\Sigma}_k = \begin{pmatrix} \sigma_k^2 & \sigma_k^2\rho_k \\ \sigma_k^2\rho_k & \sigma_k^2 \end{pmatrix}, \tag{6}$$

where $\rho_k \in (-1, 1)$ is the correlation parameter and $\sigma_k$ is the standard deviation. We further assume the $p_k$'s, $\mu_k$'s, and $\sigma_k$'s to be shared between monozygotic and dizygotic twins, with only the $\rho_k$'s being different. This is a natural assumption as marginal BMI distributions of twins of the same zygosity are expected to be identical. Therefore, we estimate $m-1$ $p_k$'s, $m$ $\mu_k$'s, $m$ $\sigma_k$'s, and $2m$ $\rho_k$'s. Furthermore, we introduce restrictions on the means to ensure identifiability of the model (See Section 4 for more details).

We select the most parsimonious number of mixture components using the criterion $\mathrm{BIC} = -2\log(L) + \log(n)Q$ for each candidate model, where $Q$ is the number of parameters and $\log(L)$ is the log likelihood function, as defined in [4, equation (3.14)].

##### 3.1 Models with a sex covariate

The mean and covariance structures (6) allow to introduce covariates easily. In the main text, we briefly described how to include a sex effect in the model in three different ways. Below, we show the mathematical formulas.

The "Stratified" model consists in estimating two completely independent Gaussian mixtures with $m$ components for male and female data, so that the parameters $\mu_k$'s, $\sigma_k$'s, $\rho_k$'s, and $p_k$'s are all sex specific. The total number of parameters is $n = 2(5m - 1)$.

The "Mean" model assumes a sex effect on $\mu_k$'s; we follow the parametrization from [4], and introduce a sex effect term $\beta_\mu$ such that for each component $k$:

$$\boldsymbol{\mu}_k = (\mu_k + \beta_\mu x_i, \mu_k + \beta_\mu x_i), \tag{7}$$

where $x_i = 0.5$ for male data and $x_i = -0.5$ for female data. Hence, $\boldsymbol{\mu}_{male} = \boldsymbol{\mu}_{female} + \beta_\mu$. The total number of parameters is $n = 5m$.

The "Mean+covariance" model assumes a sex effect on both $\mu_k$'s and $\rho_k$'s. In addition to assuming the structure 7 for $\mu_k$'s, we define the common term $\beta_\rho$ such that for each component $k$:

$$\boldsymbol{\Sigma}_k = \begin{pmatrix} \sigma_k^2 & \sigma_k^2(\rho_k + \beta_\rho x_i) \\ \sigma_k^2(\rho_k + \beta_\rho x_i) & \sigma_k^2 \end{pmatrix}, \tag{8}$$

where $x_i = 0.5$ for male data and $x_i = -0.5$ for female data. To reflect the differences between monozygotic and dizygotic correlation coefficients, we introduce two different parameters, $\beta_\rho^{MZ}$ and $\beta_\rho^{DZ}$. The total number of parameters is $n = 5m + 2$.

### 4 Implementation in TMB

To perform the analysis on the dataset, we use two files, written in two languages: R [2] and C++ [5]. We access and integrate the C++ code in the optimization process using the package TMB [6]. The code is accessible at https://github.com/skaug/Supplementary, under the repository Azzolini-etal-BMIvaries.

In the R file we read the dataset, we initialize the parameters, and we run the optimization function. This optimization function invokes the C++ script and minimizes the negative log likelihood function there defined.

The number of components of the Gaussian mixture is a hyperparameter that we select using BIC as criterion. We have already performed model selection on the dataset, so we only report the code for the best fitting model, with $m = 3$ components.

An issue that arises when Gaussian mixtures are estimated is label switching - that is, having several equivalent parameter estimates where the only difference is the order of the components. To avoid this problem, we force the means to be ordered from smallest to largest. To achieve this we reparameterize the model in terms of $\boldsymbol{\alpha}$ and we then construct the mean vector $\boldsymbol{\mu}$ as follows:

$$\mu_1 = e^{\alpha_1}$$

and, for every $2 \leq i \leq m$,

$$\mu_i = \mu_{i-1} + e^{\alpha_i}.$$

| "Stratified" | | | | | | | | | | |
|---|---|---|---|---|---|---|---|---|---|---|
| Male data | | | | | | | | | | |
| Par | $k=1$ | se | $k=2$ | se | $k=3$ | se | $k=4$ | se | Global | se |
| $\mu_k$ | 23.64 | 0.11 | 26.33 | 0.27 | 30.35 | 0.20 | | | 24.66 | |
| $\sigma_k$ | 1.97 | 0.05 | 2.46 | 0.11 | 3.67 | 0.47 | | | 2.59 | |
| $\rho_k^{(MZ)}$ | 0.72 | 0.02 | 0.27 | 0.14 | 0.24 | 0.24 | | | 0.69 | |
| $\rho_k^{(DZ)}$ | 0.34 | 0.04 | $-0.29$ | 0.12 | $-0.94$ | 0.03 | | | 0.36 | |
| $p_k$ | 0.75 | 0.05 | 0.24 | 0.05 | 0.02 | 0.01 | | | | |
| Female data | | | | | | | | | | |
| Par | $k=1$ | se | $k=2$ | se | $k=3$ | se | $k=4$ | se | Global | se |
| $\mu_k$ | 21.62 | 0.09 | 24.57 | 0.25 | 28.23 | 0.78 | | | 22.74 | |
| $\sigma_k$ | 1.91 | 0.05 | 2.83 | 0.13 | 4.91 | 0.35 | | | 2.96 | |
| $\rho_k^{(MZ)}$ | 0.75 | 0.02 | 0.34 | 0.09 | 0.41 | 0.20 | | | 0.69 | |
| $\rho_k^{(DZ)}$ | 0.26 | 0.04 | $-0.22$ | 0.08 | $-0.18$ | 0.14 | | | 0.34 | |
| $p_k$ | 0.67 | 0.04 | 0.29 | 0.03 | 0.04 | 0.01 | | | | |
| "Mean" | | | | | | | | | | |
| Par | $k=1$ | se | $k=2$ | se | $k=3$ | se | $k=4$ | se | Global | se |
| $\mu_k$ | 22.64 | 0.08 | 25.42 | 0.23 | 28.89 | 0.69 | | | 23.61 | |
| $\sigma_k$ | 1.94 | 0.04 | 2.72 | 0.12 | 4.67 | 0.32 | | | 2.84 | |
| $\rho_k^{(MZ)}$ | 0.74 | 0.02 | 0.34 | 0.08 | 0.38 | 0.15 | | | 0.69 | |
| $\rho_k^{(DZ)}$ | 0.31 | 0.03 | -0.19 | 0.07 | -0.22 | 0.12 | | | 0.36 | |
| $p_k$ | 0.70 | 0.03 | 0.26 | 0.03 | 0.04 | 0.01 | | | | |
| $\beta_\mu$ | | | | | | | | | 1.86 | 0.06 |
| "Mean+covariance" | | | | | | | | | | |
| Par | $k=1$ | se | $k=2$ | se | $k=3$ | se | $k=4$ | se | Global | se |
| $\mu_k$ | 21.41 | 0.24 | 23.17 | 0.14 | 26.15 | 0.91 | 29.68 | 0.10 | 23.57 | |
| $\sigma_k$ | 1.51 | 0.10 | 2.02 | 0.05 | 2.94 | 0.13 | 5.02 | 0.39 | 2.79 | |
| $\rho_k^{(MZ)}$ | 0.88 | 0.04 | 0.65 | 0.03 | 0.26 | 0.09 | 0.37 | 0.22 | 0.69 | |
| $\rho_k^{(DZ)}$ | 0.51 | 0.10 | 0.16 | 0.05 | -0.30 | 0.08 | -0.14 | 0.16 | 0.36 | |
| $p_k$ | 0.15 | 0.05 | 0.65 | 0.04 | 0.18 | 0.02 | 0.02 | 0.01 | | |
| $\beta_\mu$ | | | | | | | | | 1.86 | 0.06 |
| MZ $\beta_\rho$ | | | | | | | | | -0.01 | 0.03 |
| DZ $\beta_\rho$ | | | | | | | | | 0.11 | 0.05 |

Table 2: Parameter estimates for the best-fitting Gaussian mixtures for each covariate model. For each estimate, we present its standard error. The mixture components are ordered according to the value of $\sigma_k$. The global quantities, $\mu$, $\sigma$, $\rho^{(MZ)}$ and $\rho^{(DZ)}$ are calculated from [4, equation (3.4)].

| | Global quantities | | |
|---|---|---|---|
| Parameter | "Mean" | "Mean+covariance" | "Stratified" |
| male $\mu$ | 24.54 | 24.50 | 24.66 |
| female $\mu$ | 22.68 | 22.64 | 22.74 |
| male $\sigma$ | 2.84 | 2.79 | 2.59 |
| female $\sigma$ | 2.84 | 2.79 | 2.96 |
| male $\rho^{(MZ)}$ | 0.69 | 0.69 | 0.69 |
| female $\rho^{(MZ)}$ | 0.69 | 0.70 | 0.69 |
| male $\rho^{(DZ)}$ | 0.36 | 0.42 | 0.36 |
| female $\rho^{(DZ)}$ | 0.36 | 0.31 | 0.34 |

Table 3: Comparison of the global quantities for the three covariate models for both sexes.

This guarantees that $\mu_1 < \mu_2 < ... < \mu_m$. This coding works if all mean components are expected to be positive (which is the case with BMI), but can easily be tweaked so that it admits negative values as well.

In the C++ file we read the data and the parameters and we define the negative log likelihood function that is optimized in the R file. Moreover, we construct the expressions for the correlation curves and estimate them at 500 points along the BMI range. These points are used to draw the plots of the curves.

To improve the precision of our estimates, we use both the gradient and the hessian matrix in the optimization function. We are able to compute the hessian thanks to automatic differentiation performed by the package TMB.

Due to privacy, we cannot share the dataset we worked on in this document. To present the performance of the code, we created a simulated dataset. The dataset follows a Gaussian mixture distribution with three components. As parameters we used the estimates we obtained from the analysis on the twin data (Table 2). Figure 2 shows the scatterplot of the simulated data.

### 5 Comparison between different covariate models

Figure 3 is an extension of Figure 2 from the main text. The ellipses describe 0.95 probability regions for the three Gaussian components of the best-fitting model. We observe that male and female dizygotic data are both pear-shaped while that is not so evident in monozygotic data. The correlation coefficients (Table 2) well capture this difference. It also shows the higher variability of female data, which is captured by the models with a sex covariate.

Table 2 contains the parameter estimates of the best fitting Gaussian mixture within each covariate model. They also contain the standard error estimate for each coefficient.

Female data have a larger variance in its right tail (as can be seen in Figure 3) compared to male data, both for monozygotic and dizygotic twins. This is captured better by the "stratified" model. We notice that both the weights $p_2$ and $p_3$ and the standard deviations $\sigma_2$ and $\sigma_3$ are larger for female data. The lower variability in the right tail for male data, especially in dizygotic twins, is also reflected in the correlation coefficient $\rho_3^{(DZ)}$ which approaches $-1$, compared to the value of $-0.18$ for female data. Despite these differences being justified by the dataset, the BIC value still prefers the less flexible mean covariate model.

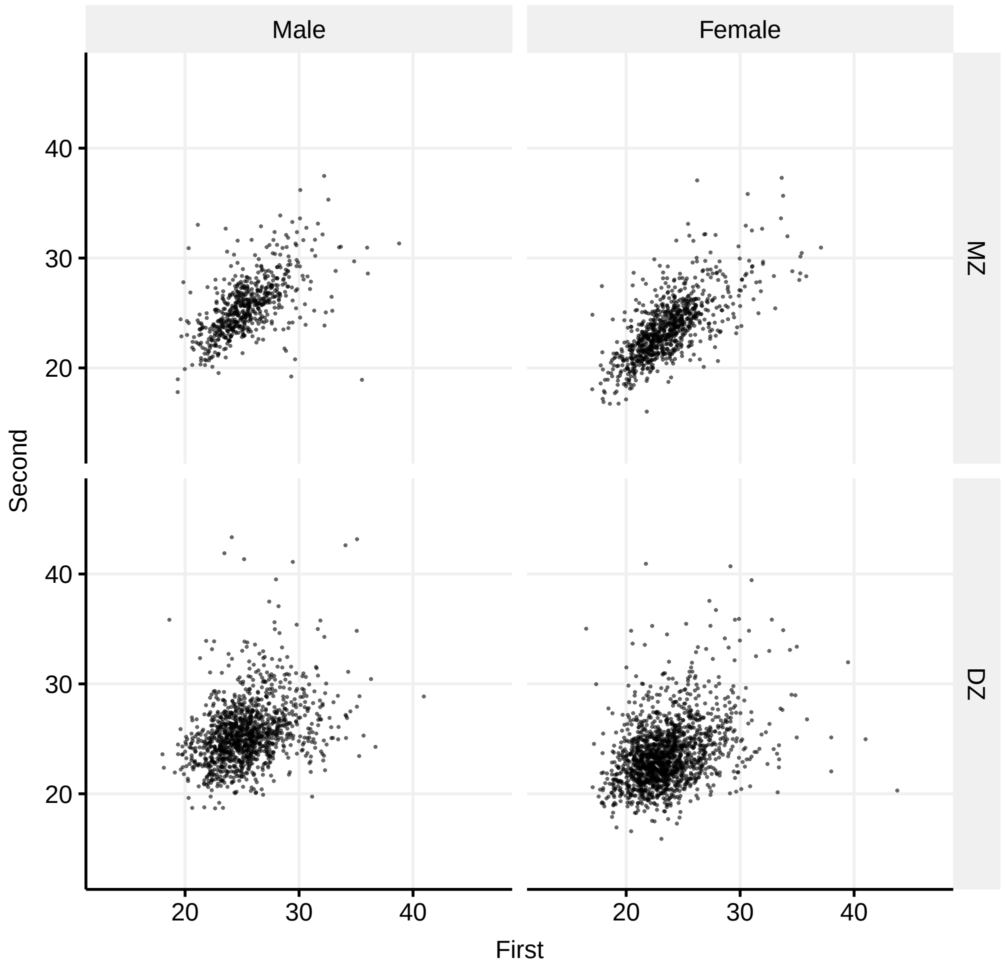


Figure 2: Scatterplot of the simulated dataset, divided by sex (male and female) and zygosity (monozygotic and dizygotic). The dataset follows a Gaussian mixture distribution, using as parameters the estimates from our analysis.

The values of $\mu_k^F$ and $\mu_k^M$ obtained through the stratified analysis are quite similar to the ones from the best fitting model (mean covariate), albeit slightly larger. The average difference between male and female mean components for stratified analysis is 1.97, slightly larger than the estimated parameter $\beta_\mu = 1.86$. This is not reflected in the global quantities (Table 3), which are very similar between the two different models.

Comparing the "mean+covariance" parameter estimates (Table 2) is slightly more difficult, since the best fitting mixture has one component more than the other two best fitting models. Looking at the global quantities (Table 3), we see that they are pretty consistent with the other two models. Moreover, the coefficient $\beta_\mu$ is the same as in the "Mean" model.

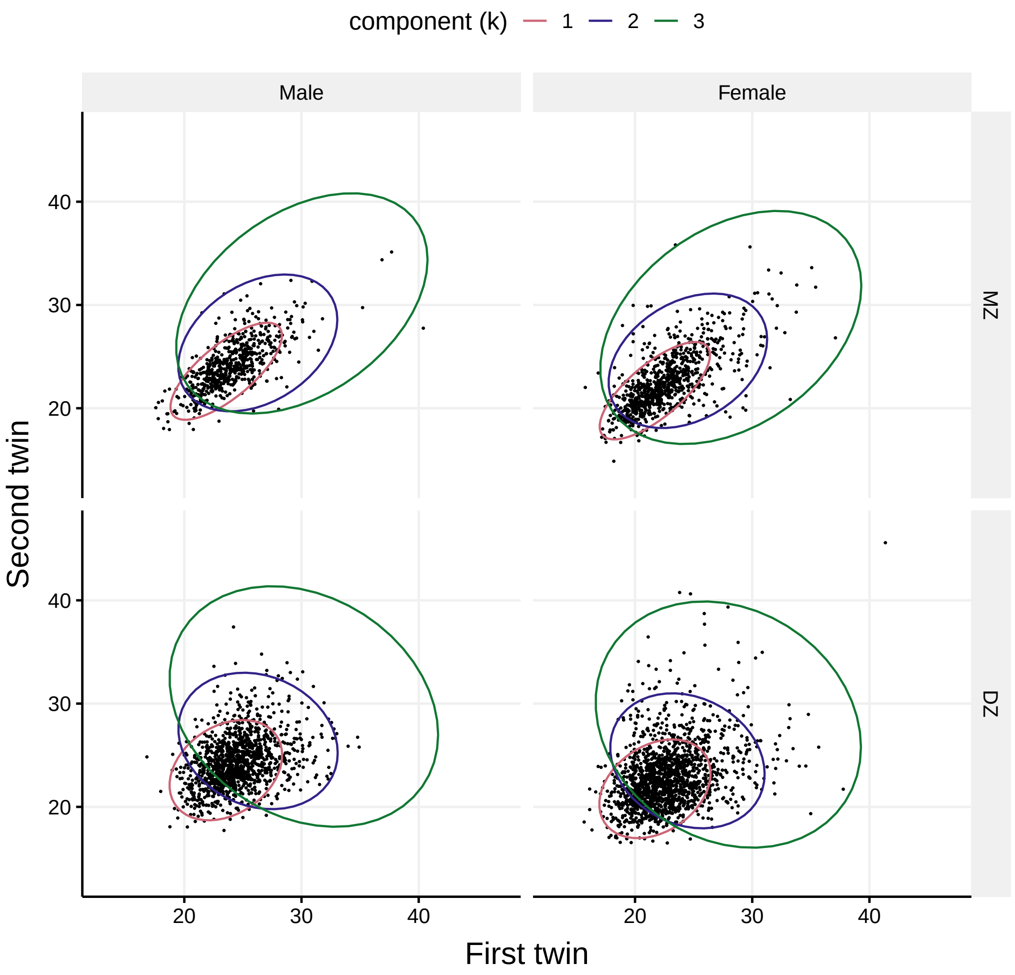


Figure 3: Ellipses representing the three Gaussian kernels, divided by sex and zygosity.

The coefficient $\beta_\rho^{(MZ)}$ is not significant. The model does identify a more significant sex effect on the dizygotic correlation coefficient. The BIC values still favors the simpler "Mean" model.

#### 5.1 Comparison of correlation curves

The differences in BIC values between the best-fitting mixtures among different covariate models is not very large (see Table 1 in main text). We also showed, in the above section, that the parameter estimates are relatively consistent between the different models. In this section we compare the monozygotic correlation curves obtained using the models from the previous section ("stratified", "Mean", "Mean+Covariance").

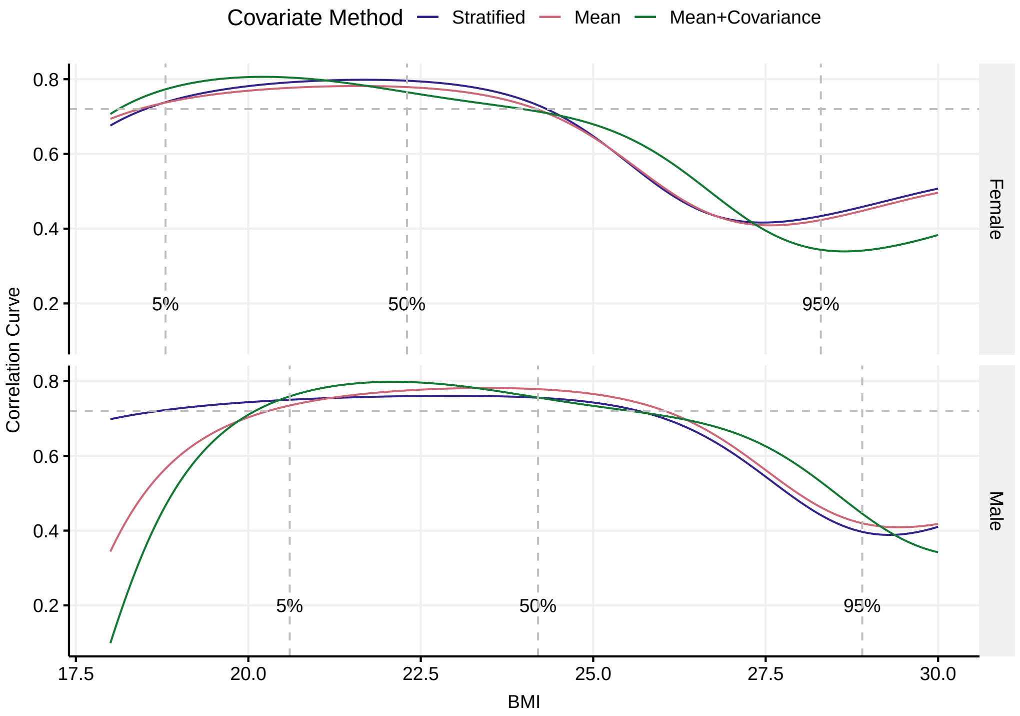


Figure 4: Correlation curves for monozygotic twins using the three different methods described in section 3.1. For each method, we plotted the correlation curve for the best model for both male and female BMI. The horizontal grey lines represent the Pearson global correlation $\rho^{(MZ)}$. The vertical dotted lines represent the 0.05, 0.5, and 0.95 quantiles of the data, divided by sex.

Figure 4 displays the estimated monozygotic correlation curves according to the different models. For each of the three methods, we plot a curve for male data and one for female data. The three vertical lines represent the 0.05, 0.50, and 0.95 quantiles of the male and female data separately.

We observe that the curves of the same sex follow the same broad shape. In general, "stratified" and "mean" models generate curves that are quite similar. We remark that, while the "mean" model generates male and female curves which are identical and shifted on the x-axis, the same is not true for the more flexible "stratified" model. This is particularly evident by looking at the distance between male and female curves in the left tail.

The curves generated by the "mean+covariance" model deviate slightly from the pattern above, especially in the right tail. This difference is possibly caused by the different number of components of the best fitting model.

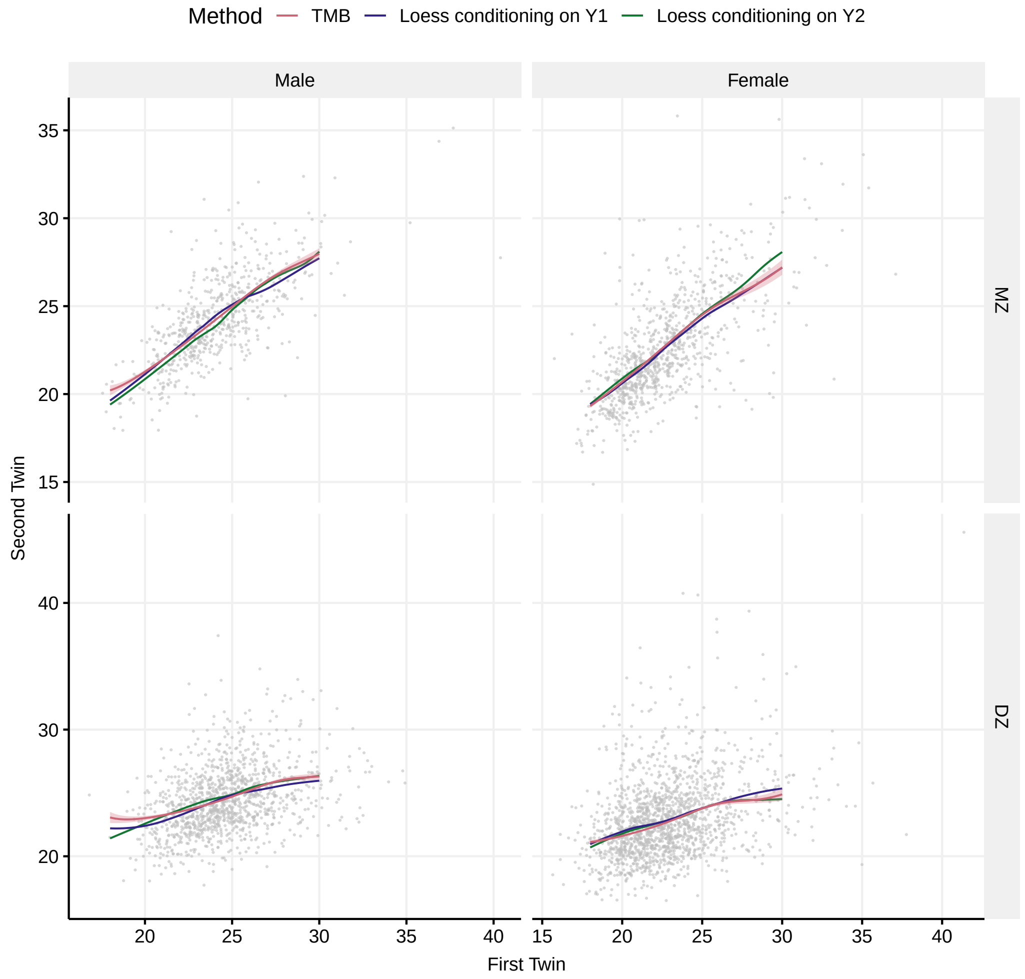


Figure 5: Comparison of estimated conditional mean curves based on the Gaussian mixture (TMB; red) with the two fully nonparametric estimates (blue and green) obtained from "loess". The estimation uncertainty is displayed only for the Gaussian mixture, and the underlying data are shown as grey dots.

### 6 Sensitivity Study

As a sensitivity study, we compared the estimated correlation curve based on the Gaussian mixture with a fully nonparametric method based on the smoother function "loess" in R. In particular, we computed nonparametric conditional means and variances, and these were used to obtain nonparametric correlation curves. In "loess" we first regressed $Y1$ on $Y2$, and secondly $Y2$ on $Y1$, and hence obtained two different estimated correlation curves. Due to the arbitrary labeling of twin pair members these curves are estimates of the same quantity,

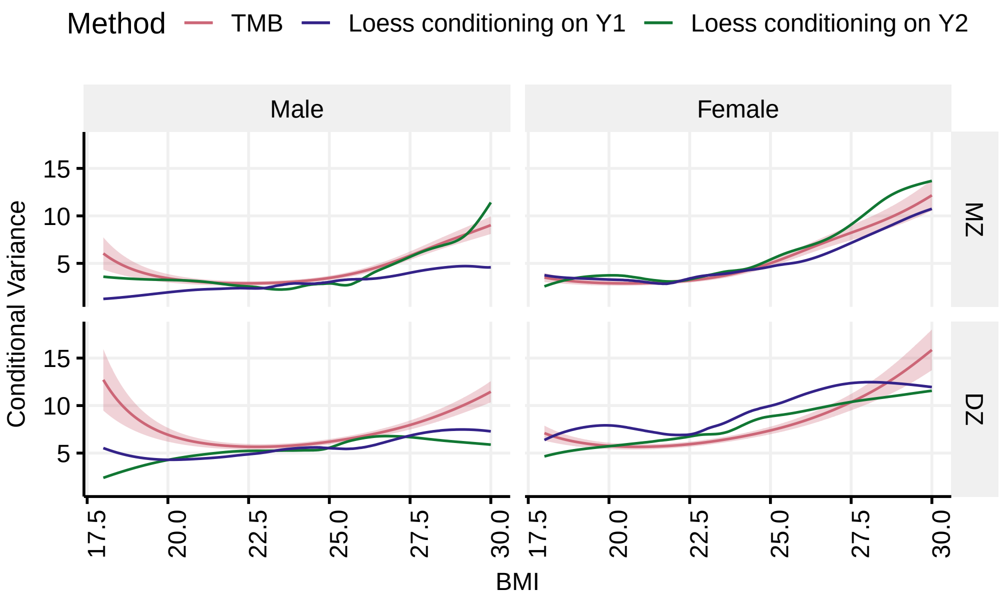


Figure 6: Comparison of estimated conditional variance curves based on the Gaussian mixture (TMB; red) with the two fully nonparametric estimates (blue and green) obtained from "loess". The estimation uncertainty is displayed only for the Gaussian mixture.

and differ only due to estimation uncertainty. As we are estimating a conditional mean curve, we will refer to "regressing $Y1$ on $Y2$" as "conditioning on $Y2$". We show a simplified version of the code in appendix B.

Looking at Figure 5, 6 and 7 we see that, overall, female curves among different models seem more similar to each other compared to those of males. A similar observation can be made for dizygotic curves compared monozygotic ones. We remark that the sample sizes of the subsets are quite different; there are about 30% more female subjects than male subjects, and the size of the dizygotic subset is double the size of the monozygotic subset. The tail behaviour is sometimes discordant with the previous remark, especially in Figure 7. We investigate this phenomenon later.

Figure 5 shows the conditional means obtained using TMB and "loess" conditioning on both twin pairs separately. The three curves identify the same pattern in the data, although they do not coincide perfectly along the entire BMI range. We observe a larger variance in the tails, which can be expected, given the sparsity of the dataset.

The two nonparametric conditional variance curves lie partly outside the pointwise 95% confidence intervals for the Gaussian mixture (Figure 6), particularly in the tails. There is also a discrepancy between the two nonparametric curves, and this discrepancy can be interpreted as estimation uncertainty, because the blue and green curves are the same estimator applied to two different datasets (switching the roles of $Y1$ on $Y2$).

To compute the derivative of the conditional mean curve needed for the correlation curve,

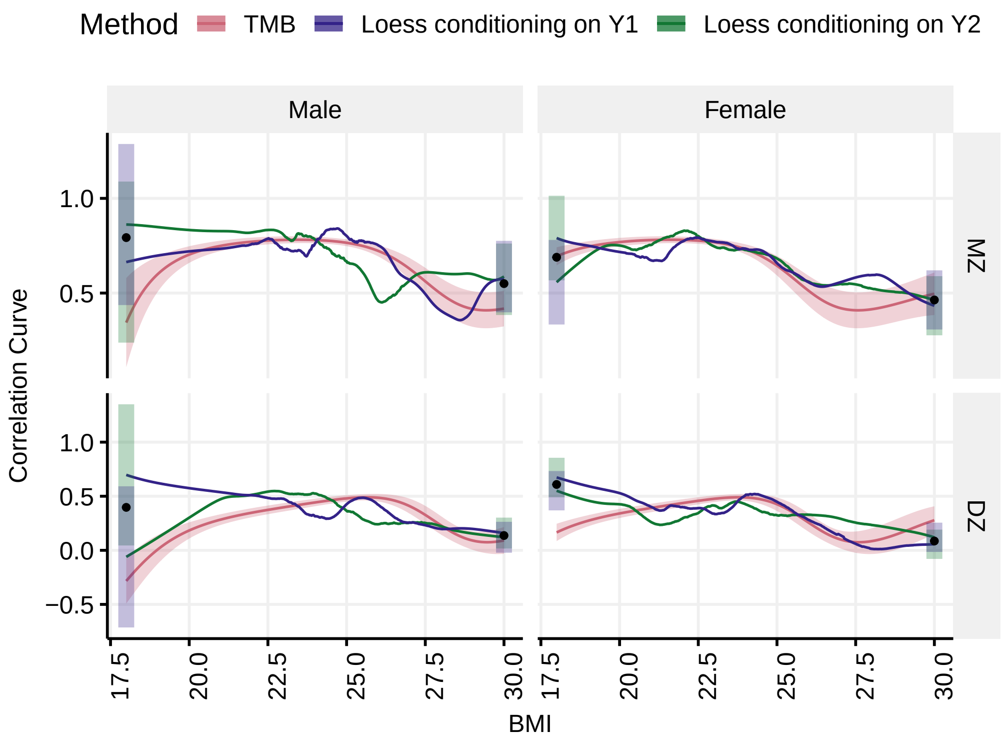


Figure 7: Comparison of estimated correlation curves based on the Gaussian mixture (TMB; red) with the two fully nonparametric estimates (blue and green) obtained from "loess". For the Gaussian mixture the estimation uncertainty is displayed for the full range of BMI values, while for the two nonparametric curves 95% confidence intervals are displayed only for BMI=18 and 30. The black dot represents the average of the pooled 100 bootstrap samples.

we used the finite difference approximation

$$\beta(y) = \frac{\mu(y+h) - \mu(y-h)}{2h}, \tag{9}$$

for $h = 0.1$, where $\mu(y)$ is the conditional mean obtained from "loess". In Figure 7 we show the loess estimate of the correlation curve. Overall, they show the same results as the TMB estimate. The behaviour of the curves differ mostly in the tails. We investigate this, and in particular the left-hand tail for dizygotic females, in more detail below.

To this end we generated 100 bootstrap samples from the dataset, sampling twin pairs with replacement and randomly reassigning the order of the twin members. For each of the 100 bootstrap datasets we computed the nonparametric correlation curve, and subsequently calculated bootstrap mean and standard deviation. Since the largest difference between correlation curves is in the tails, Figure 7 displays the bootstrap results only for BMI=18 and

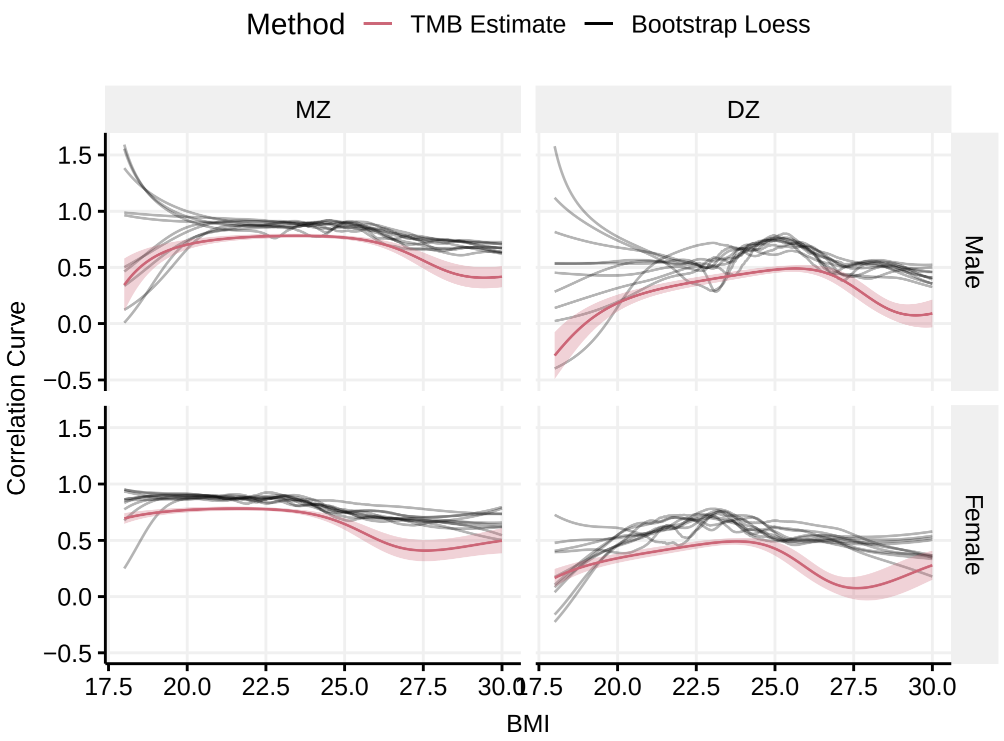


Figure 8: Fully nonparametric correlation curves estimated on simulated data (in black) compared to the parametric TMB estimate (in red). The estimation uncertainty is displayed only for the Gaussian mixture.

30. The 95% bootstrap confidence intervals are constructed using 1.96 standard deviations around the curve.

The confidence intervals for male data at BMI=18 are very wide; this is probably a reflection of the few data points around that value in the original dataset. Looking at the female dizygotic data at BMI=18, neither confidence intervals (blue or green) overlap with the TMB estimate. Hence, the differences cannot be described by uncertainty alone and may be attributable to either lack of flexibility of the Gaussian mixture (TMB) or to some bias in the nonparametric estimator.

To investigate the statistical properties of the nonparametric estimator, we conducted a simulation experiment in which data were simulated from the fitted Gaussian mixture (based on the real data). Because an analytic expression for the correlation curve is available for Gaussian mixtures [4] we can compare the nonparametric estimator with the "true" correlation curve. Figure 8 shows that the nonparametric estimator has an upwards bias throughout the BMI range. Strictly speaking, this conclusion is valid only when the true data generating mechanism is a Gaussian mixture, but it seems necessary to explore the properties of the nonparametric estimator further.

### A Data preprocessing, R code

Below, we show a simplified version of the preprocessing code and we apply it to a dummy dataset. The dataset contains four measurements of two twins belonging to the same pair.

```
 Dataset
##   Tvparnr Twinnumber Twinid Wave Age Sex Zygoti  BMI
## 1       1          1     11    1  20   1      1 21.3
## 2       1          1     11    2  30   1      1 22.5
## 3       1          1     11    3  37   1      1 23.2
## 4       1          1     11    4  47   1      1 25.1
## 5       1          2     12    1  20   1      1 21.6
## 6       1          2     12    2  30   1      1 22.4
## 7       1          2     12    3  37   1      1 22.7
## 8       1          2     12    4  47   1      1 23.6
library(tidyverse)
value<-c()
newdat<-data.frame(Age=35)
IDs<-unique(Dataset$Twinid)
for(j in IDs) {
   Subset<-Dataset%>%filter(Twinid==j)
   if(dim(Subset)[1]>2){
   lm_mod<-lm(BMI~Age, data = Subset)
   }
 P<-predict(lm_mod, newdata = newdat)
 value<-c(value, P)
}

value
       1        1
## 23.23226 22.68316
```

The code separates the dataset depending on the variable "Twinid", checks if the number of waves is larger than two (to reduce uncertainty we only consider twin pairs with three or more measurements), and performs a linear regression on said dataset. The output "value" is the predicted BMI value at age 35 and forms the dataset used in the analysis.

### B Nonparametric correlation curve, R code

The code below is a simplified version of the code used to compute the nonparametric correlation curve. Assume that the dataset called "Dataset" contains data from one single zygosity and one sex (for example, only male monozygotic twins). The code shows how we compute conditional variance and the derivative of the conditional mean using "loess". We only condition on $Y2$, since conditioning on $Y1$ follows the same procedure, only with the role of $Y1$ and $Y2$ switched.

The quantity "Fit-square" corresponds to $\mathbb{E}(Y1^2 \mid Y2)$. To compute the conditional variance, we then calculate $\mathbb{E}(Y1^2 \mid Y2) - \mathbb{E}(Y1 \mid Y2)^2$.

To compute the derivative of the conditional mean, we use equation 9 for $h = 0.1$.

```
library(tidyverse)
vec<-seq(18, 30, length.out =500)
newdata<-data.frame(First=vec,Second=vec,
                    Firstsq=vec*vec, Secondsq=vec*vec)
new_loess<-function(data, formula){
      loess(formula, data,
               control=loess.control(surface="direct"))
}
## conditional mean
CM<-Dataset%>%new_loess(First~Second)%>%
  predict(newdata=newdata)
## conditional variance
Fit_square<-Dataset%>%new_loess(Firstsq~Second)%>%
  predict(newdata=newdata)
CV<-Fit_square-CM*CM
## derivative of the conditional mean (beta)
h<-0.1
newdata_plus<-newdata+h
newdata_minus<-newdata-h
Fit_plus<-Dataset%>%new_loess(First ~Second)%>%
  predict(newdata=newdata_plus)
Fit_minus<-Dataset%>%new_loess(First ~Second)%>%
  predict(newdata=newdata_minus)
Beta<-(Fit_plus-Fit_minus)/(2*h)
## correlation curve
Sigma_Beta<-sd(Dataset$Second)*Beta
Sigma_Beta_squared<-Sigma_Beta*Sigma_Beta
Correlation_curve<-Sigma_Beta/sqrt(Sigma_Beta_squared+CV)
```

[3] CE McCulloch and JM Neuhaus. *Generalized linear mixed models*. Wiley Online Library, 2001.

[4] G D Berentsen, F Azzolini, H J Skaug, R T Lie, and H K Gjessing. Heritability curves: A local measure of heritability in family models. *Statistics in Medicine*, 2020.

[5] ISO/IEC. Programming languages — c++. Draft International Standard N4660, March 2017. URL https://web.archive.org/web/20170325025026/http://www.open-std.org/jtc1/sc22/wg21/docs/papers/2017/n4660.pdf.

[6] K Kristensen, A Nielsen, CW Berg, HJ Skaug, and BM Bell. TMB: Automatic differentiation and Laplace approximation. *Journal of Statistical Software*, 70(5):1–21, 2016. doi: 10.18637/jss.v070.i05.
